# Mitotic chromosome alignment is required for proper nuclear envelope reassembly

**DOI:** 10.1101/343475

**Authors:** Cindy L. Fonseca, Heidi L.H. Malaby, Leslie A. Sepaniac, Whitney Martin, Candice Byers, Anne Czechanski, Dana Messinger, Mary Tang, Ryoma Ohi, Laura G. Reinholdt, Jason Stumpff

## Abstract

Chromosome alignment at the equator of the mitotic spindle is a highly conserved step during cell division, however, its importance to genomic stability and cellular fitness are not understood. Normal mammalian somatic cells lacking Kif18A function complete cell division without aligning chromosomes. These alignment-deficient cells display normal chromosome copy numbers *in vitro* and *in vivo*, suggesting that chromosome alignment is largely dispensable for maintenance of euploidy. However, we find that loss of chromosome alignment leads to interchromosomal compaction defects during anaphase, abnormal organization of chromosomes into a single nucleus at mitotic exit, and the formation of micronuclei *in vitro* and *in vivo*. These defects slow cell proliferation and reduce postnatal growth and survival with variable penetrance in mice. Our studies support a model in which the alignment of mitotic chromosomes promotes proper nuclear envelope reassembly and continued proliferation by ensuring that chromosomes segregate as a compact mass during anaphase.

## INTRODUCTION

Chromosome alignment at the mitotic spindle equator is a conserved feature of cell division in eukaryotic cells, suggesting that it has an essential function for accurate chromosome segregation. Possible functions of chromosome alignment include promoting attachments between chromosomes and spindle microtubules, preventing erroneous attachments, promoting equal chromosome segregation during anaphase, and coordinating anaphase and cytokinesis (Kops et al., 2010; Maiato et al., 2017). Elucidating the importance of chromosome alignment has been technically difficult due to an inability to experimentally disrupt chromosome alignment without also altering attachments between kinetochores and spindle microtubules. Thus, it remains unclear how chromosome misalignment per se contributes to defects in chromosome copy number, development, and disease. New experimental models are, therefore, needed to address the functional importance of chromosome alignment to cellular and organismal physiology.

In mammalian cells, metaphase alignment requires the confinement of bioriented chromosome pairs to the spindle equator region. While the majority of chromosome pairs are located near the center of the spindle at the start of mitosis, some must be transported to the equator through a process called congression (Kapoor et al., 2006; Magidson et al., 2011). Paired chromosomes establish end-on attachments to microtubules emanating from opposite spindle poles via kinetochores, which assemble at the centromeric region of each chromosome. These bioriented chromosomes undergo microtubule-driven, oscillatory movements that initially permit excursions away from the equator (Skibbens et al., 1993). Therefore, the alignment of bioriented chromosomes requires the regulation of kinetochore-attached microtubules in a way that dampens these oscillations and limits them to an area around the spindle center.

Congression, biorientation, and chromosome confinement rely on kinesin-dependent mechanisms. CENP-E (kinesin-7) transports mono-oriented chromosomes to the spindle equator and works synergistically with KIF22 (kinesin-10) to promote the biorientation of chromosome pairs (Barisic et al., 2014; Drpic et al., 2015; Kapoor et al., 2006; Schaar et al., 1997). Loss of CENP-E or KIF22 function leads to chromosome segregation defects both *in vitro* and *in vivo* (Ohsugi et al., 2008; Weaver et al., 2003). However, the majority of chromosomes are able to align in cells lacking either CENP-E or KIF22 (Levesque and Compton, 2001; Putkey et al., 2002; Schaar et al., 1997), and the presence of attachment defects under these conditions complicates determination of the primary defect underlying chromosome segregation errors. Another kinesin motor, KIF18A (kinesin-8), is primarily responsible for the confinement of chromosome movements during metaphase (Mayr et al., 2007; Zhu et al., 2005). KIF18A concentrates at the plus-ends of kinetochore microtubules and functions to reduce chromosome movements through direct suppression of kinetochore-microtubule dynamics (Stumpff et al., 2008; 2012). Therefore, loss of KIF18A disrupts the alignment of all chromosomes.

Unlike CENP-E and KIF22, a role for KIF18A in promoting proper kinetochore-microtubule attachments is cell type specific. Germ cells, as well as some genomically unstable tumor cell lines, require KIF18A function to satisfy the spindle assembly checkpoint and promote the metaphase to anaphase transition (Czechanski et al., 2015; Mayr et al., 2007; Zhu et al., 2005). These data suggest KIF18A has a role in establishing or maintaining kinetochore microtubule attachments. In contrast, primary mouse embryonic fibroblasts lacking KIF18A function progress through mitosis with normal timing, despite failing to align chromosomes (Czechanski et al., 2015). Thus, KIF18A’s alignment and attachment functions appear to be separable. Accordingly, *Kif18A* mutant mice survive to adulthood, although at slightly lower than the expected Mendelian ratio (Czechanski et al., 2015; Reinholdt et al., 2006). Collectively, these data implicate KIF18A-deficient somatic cells as a useful model system to determine the consequences of division with unaligned, but correctly attached, chromosomes.

Here, we show that mitotic cell division in the absence of chromosome alignment does not significantly alter chromosome copy number. Instead, chromosome alignment is required for proper nuclear envelope reformation and the organization of all chromosomes into a single nucleus. These defects reduce proliferation, leading to growth defects and sub-viability in mice. Our results define the physiological role of chromosome alignment independent of chromosome attachment, highlighting the importance of metaphase chromosome organization for proper nuclear envelope templating and proliferation in the next cellular generation.

## RESULTS

### Human somatic cells deficient for KIF18A function divide in the absence of chromosome alignment

To determine if chromosome alignment is required for cell division in normal human cells, we tested the effects of KIF18A depletion on mitotic chromosome organization and progression through mitosis in a human retinal pigment epithelial cell line (RPE1) immortalized by human telomerase expression (hTERT). hTERT-RPE1 cells were derived from a female and are near diploid, containing a modal chromosome number of 46 with a single derivative X-chromosome. These cells have a robust spindle assembly checkpoint response and display chromosome segregation errors under conditions that promote abnormal kinetochore microtubule attachments (Thompson and Compton, 2008; Uetake and Sluder, 2010).

To determine the effects of KIF18A depletion on chromosome alignment in normal human somatic cells, we treated hTERT-RPE1 cells with control or *KIF18A*-specific siRNAs and then fixed and stained for kinetochores and centrosomes. We quantified chromosome alignment by measuring the distribution of kinetochores along the spindle axis, as previously described (Fonseca and Stumpff, 2016; Stumpff et al., 2012). The distribution of kinetochores within the spindle was significantly increased in hTERT-RPE1 cells 48, 96, and 144 hours after *KIF18A* knockdown (KD) (Figure 1 A-B). To achieve complete ablation of KIF18A function, we used CRISPR-Cas9 genomic editing to generate a homozygous *KIF18A* knockout (KO) hTERTRPE1 line. Using antibodies that recognize either the C-terminus or the motor domain of KIF18A, we confirmed that these *KIF18A*-KO cells have undetectable KIF18A protein. (Figure S1). We found that the null cells also display chromosome alignment defects similar to those seen in *KIF18A*-KD cells. Importantly, expression of EGFP-KIF18A, but not EGFP alone, rescues chromosome alignment in *KIF18A*-KO cells (Figure 1 C-D). Thus, the chromosome alignment defects in *KIF18A*-KO cells are due specifically to the loss of KIF18A activity.

**Figure 1.**
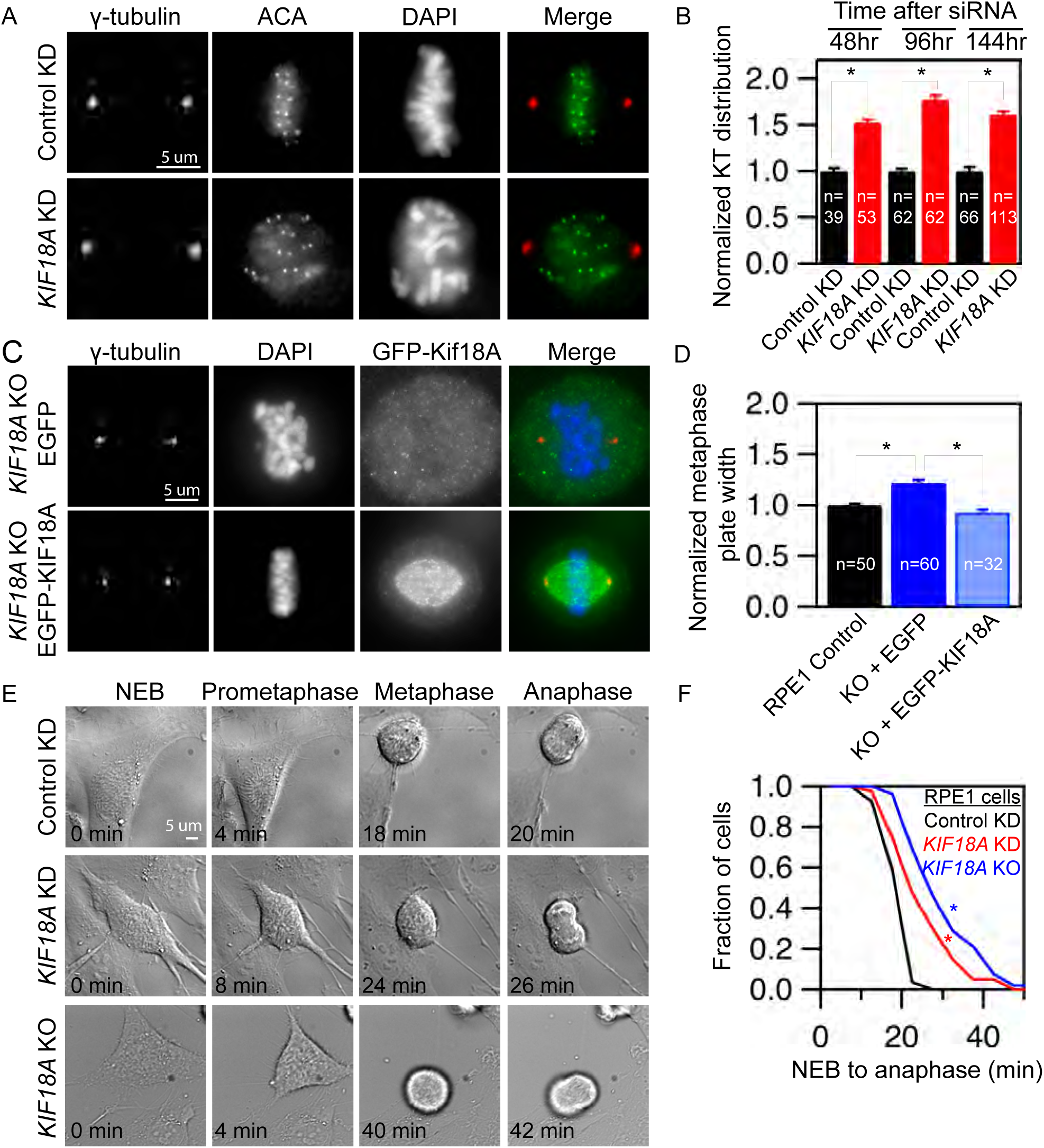
Human Retinal pigment epithelial cells lacking KIF18A function progress through mitosis with unaligned chromosomes. (A) Representative images of centrosomes and kinetochores in fixed hTERT-RPE1 cells treated with the indicated siRNAs. (B) Plot of kinetochore (KT) distribution at the indicated times following siRNA treatment measured using the full-width at half maximum of kinetochore fluorescence along the pole-to-pole axis. (C) Images of *KIF18A* knockout (“KO”) cells transiently expressing EGFP or EGFP-KIF18A. Cells were fixed and stained for γ-tubulin and DNA. (D) Plot of metaphase plate width in control and *Kif18A* KO cells expressing GFP or GFP-KIF18A. Plate width was determined by measuring the full width at half maximum of DAPI fluorescence along the pole-to-pole axis. (E) Stills from time-lapse DIC imaging of hTERT-RPE1 cells from the indicated treatment groups. (F) Cumulative frequency plot of time from nuclear envelope breakdown (NEB) to anaphase onset. n= 27 (control), n= 40 (*KIF18A* KD), and n = 52 (*KIF18A* KO). Data sets were compared using a Kruskal-Wallis one-way ANOVA with Dunn’s multiple comparisons test, asterisks (*) indicate p<0.01, error bars represent SEM.

We used live cell imaging to directly determine whether hTERT-RPE1 cells require KIF18A for progression through mitosis. In contrast to the long, spindle assembly checkpoint dependent mitotic arrest displayed by KIF18A-depleted HeLa cells (Mayr et al., 2007; Stumpff et al., 2008; Zhu et al., 2005), the time from nuclear envelope breakdown to anaphase is moderately extended from a mean of 20.0 ± 3.3 minutes in control cells to 25.6 ± 8.2 and 31.0 ± 10.5 minutes in *KIF18A*-KD and KO hTERT-RPE1 cells, respectively (Figure 1 E-F). Taken together, these data indicate that hTERT-RPE1 cells lacking KIF18A function are able to complete cell division in the absence of chromosome alignment, similar to primary embryonic fibroblasts derived from *Kif18A^gcd2/gcd2^* mutant mice, which carry a mutation that completely inactivates KIF18A (Czechanski et al., 2015). These features permitted us to use *KIF18A* loss of function cells from mouse and human as tools to determine the consequences of undergoing cell division without mitotic chromosome alignment.

### Chromosome alignment is largely dispensable for equal chromosome segregation

To determine whether chromosome alignment is required to maintain proper chromosome copy number, we analyzed hTERT-RPE1 cells treated with *KIF18A* siRNAs for 6 days using chromosome specific fluorescence *in situ* hybridization (FISH) probes. hTERT-RPE1 cells display a doubling time of 14-24 hours (Uetake and Sluder, 2004), and therefore, are expected to complete approximately 6 divisions in the absence of chromosome alignment during the treatment period. KIF18A knockdown did not significantly alter the copy number of the 10 chromosomes we analyzed when compared to control siRNA treated cells (Figure 2 AB, Table S1). As a positive control, we analyzed chromosome copy number in hTERT-RPE1 cells treated with siRNAs targeting the spindle assembly checkpoint protein MAD2. Depletion of MAD2 promotes premature anaphase chromosome segregation prior to chromosome alignment in the presence of abnormal kinetochore microtubule attachments, which lead to chromosome segregation errors (Canman et al., 2002; Meraldi et al., 2004; Michel et al., 2001). In contrast to *KIF18A* knockdown cells, *MAD2* knockdown cells were aneuploid for all chromosomes tested (Figure 2B, Table S1), consistent with previous reports (Michel et al., 2001). We also measured the total number of chromosomes present in mitotic *KIF18A*-KD and *KIF18A*-KO RPE1 cells, as well as primary embryonic fibroblasts from *Kif18A^gcd2/gcd2^* mice. In all cases, chromosome numbers were comparable to those found in matched control cells (Figure 2 C-D).

**Figure 2.**
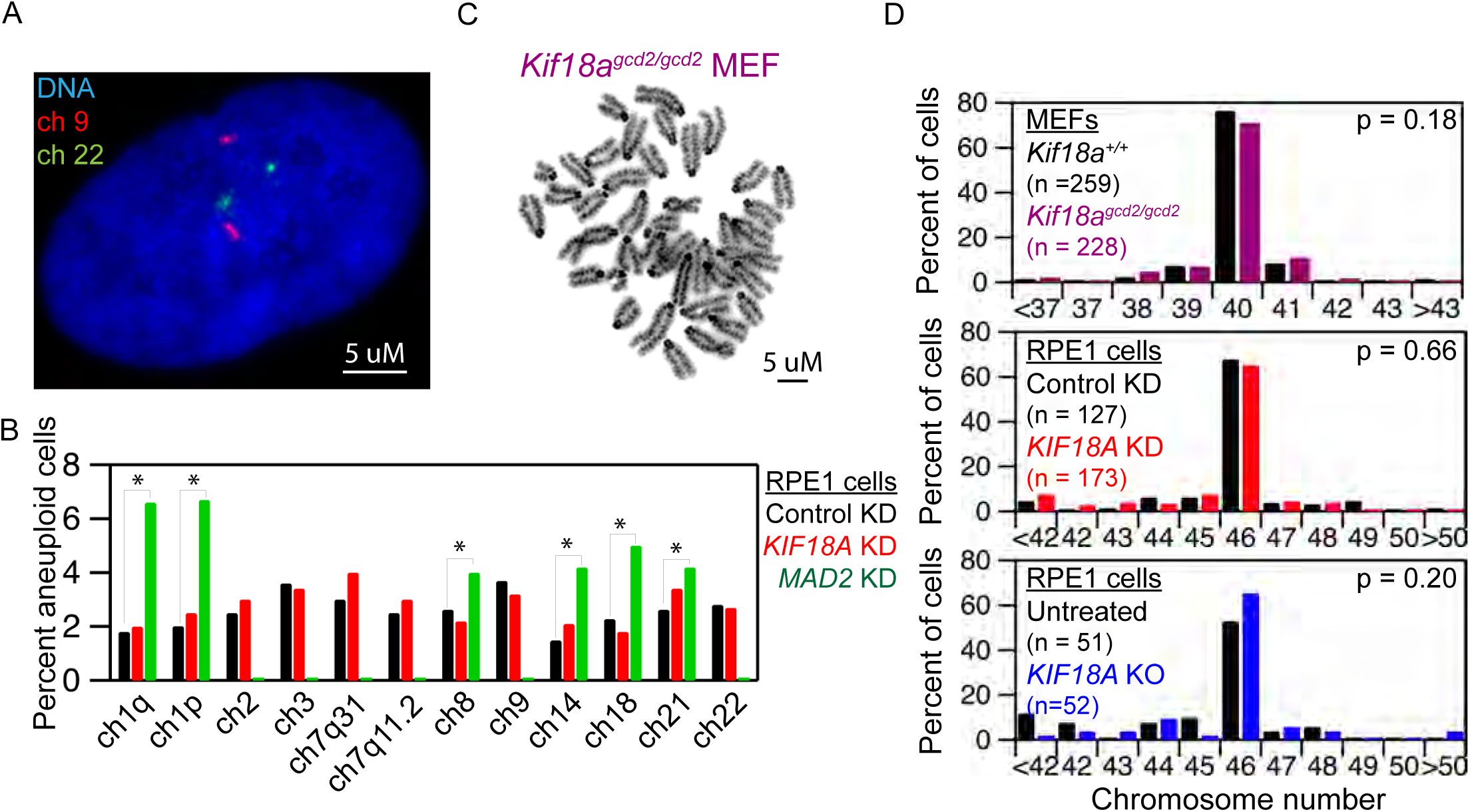
Loss of KIF18A function does not alter chromosome copy number. (A) Image of an hTERT-RPE1 cell with chromosomes 9 (red) and 22 (green) labeled by fluorescent *in situ* hybridization (FISH). (B) Plot of the percentage of cells aneuploid for the indicated chromosomes following treatment with control siRNA, *KIF18A* siRNA, or *MAD2* siRNA. N = 1500 cells for each condition, asterisks (*) indicate p<0.05 based on Chi-Squared analyses. The effect of *MAD2* KD on chromosomes 2, 3, 7, or 9 was not determined. (C) Image of Geimsa stained metaphase spread of an early passage (p0), mouse embryonic fibroblast (MEF). (D) Quantification of metaphase chromosome numbers from (top) wild type and *Kif18a^gcd2/gcd2^*, early passage (p0), pair-matched, MEFs; (middle) hTERT-RPE1 cells treated with control or KIF18A siRNAs; and (bottom) control and KIF18A KO RPE1 cells. Indicated p-values were calculated by Chi-Squared analyses.

To test the possibility that aneuploid KIF18A-deficient cells were being eliminated from the population via apoptosis, we measured the percentage of *KIF18A*-KD cells positive for cleaved caspase-3, an apoptotic marker. *KIF18A*-KD cells did not display an increase in cleaved caspase-3 relative to control treated cells 48 or 144 hours after siRNA treatment (Figure S2). Furthermore, our previous analyses of apoptosis in MEFs from *Kif18A^gcd2/gcd2^* mice indicated no increase in TUNEL-positive cells compared to wild type MEFs (Czechanski et al., 2015). Taken together, these data indicate that chromosome alignment per se is largely dispensable for the maintenance of euploidy and that abnormal kinetochore microtubule attachments likely underlie segregation errors in *MAD2*-depleted cells.

### Loss of KIF18A leads to nuclear organization defects in interphase cells

Despite the lack of evidence for chromosome copy number changes, we observed that KIF18A-deficient interphase cells display defects in nuclear organization. Loss of KIF18A function leads to an increase in cells with micronuclei (Figure 3A). The percentage of *KIF18A*KD hTERT-RPE1 cells with micronuclei increases with time following siRNA treatment (Figure 3B). Similarly, both *KIF18A*-KO hTERT-RPE1 cells and *Kif18A^gcd2/gcd2^* MEFs display a significant increase in micronuclei compared to control cells (Figure 3 C-D). The majority of these micronuclei (~70-85%) are positive for centromeres, consistent with most containing whole chromosomes (Figure 3E). Kif18A deficiency also leads to the formation of micronuclei *in vivo*. The percentage of micronucleated reticulocytes is significantly increased in both *Kif18A^gcd2/gcd2^* and in *Kif18A^gcd2/+^* mice relative to controls in an additive manner (Figure 3F). The level of micronucleated reticulocytes in *Kif18A^gcd2/gcd2^* mice is comparable to that of mice with genomic instability due to deficient DNA double strand break repair (ataxia telangiectasia-mutated, *ATM^tm1Awb/tm1Awb^*) (Figure 3F). These results indicate that KIF18Adependent chromosome alignment is required to prevent micronucleus formation, and that a single functional allele of *Kif18a* is not sufficient for normal genetic stability.

**Figure 3.**
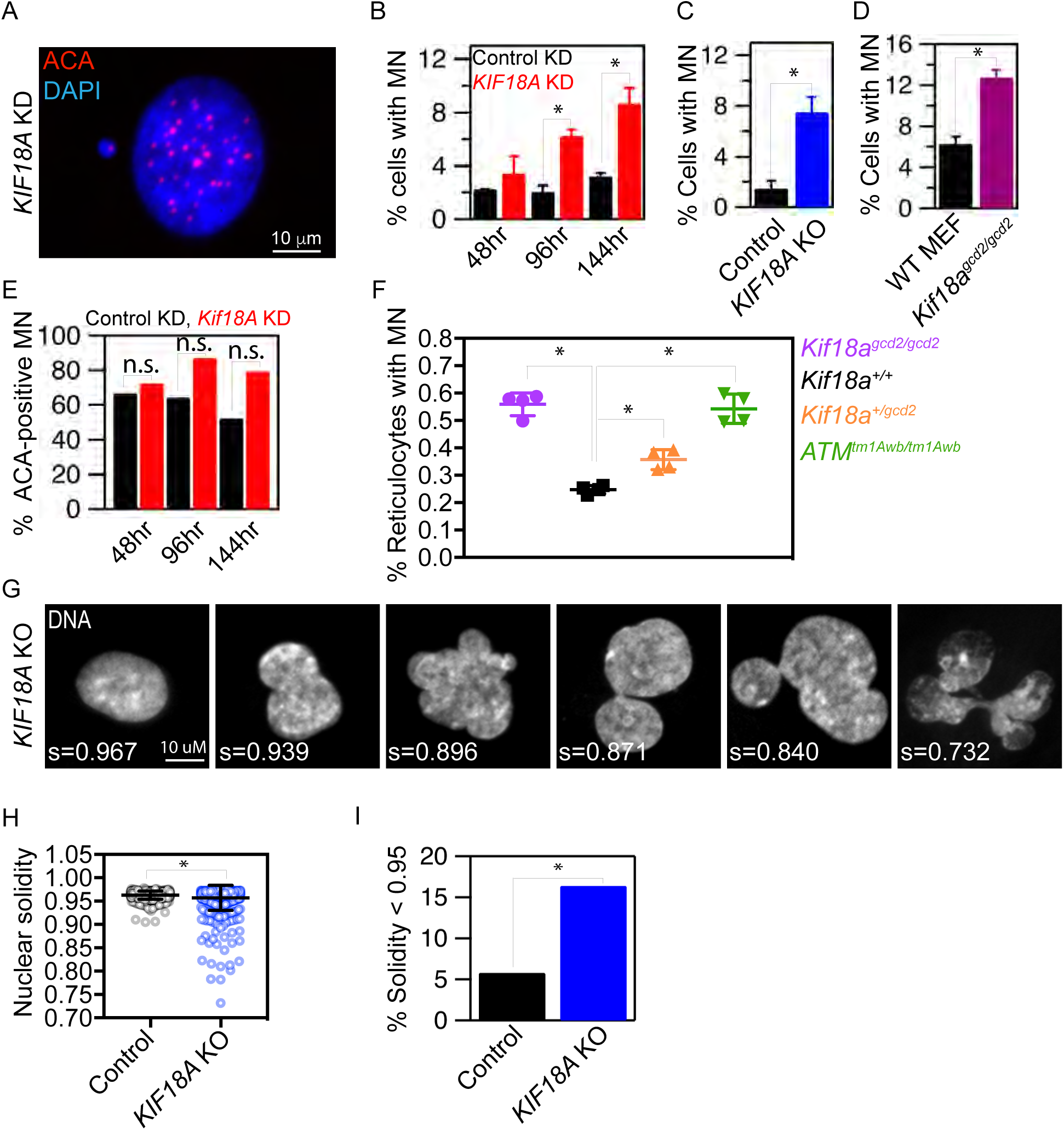
KIF18A-deficient cells form micronuclei and abnormal nuclear shapes. (A) Representative image of a micronucleated hTERT-RPE1 cell labeled with DAPI and anticentromere antibodies (ACA) to visualize DNA and centromeres, respectively. (B-D) Plots of the percentage of cells with micronuclei (MN) in (B) cells treated with the indicated siRNAs for 48h, 96h, or 144h; (C) control and *KIF18A* KO hTERT-RPE1 cells; and (D) mouse embryonic fibroblasts (MEF) from *Kif18a^gcd2/gcd2^* mice. n > 600 cells for each condition, data were compared via Chi-Squared analyses. (E) Quantification of the percentage of micronuclei containing centromeric DNA (ACA-positive) in control (black) and *KIF18A* siRNA (red) treated cells. (F) Quantification of micronuclei in mouse peripheral blood reticulocytes from *Kif18a^+/+^*, *Kif18a^gcd2/gcd2^*, *Kif18a^+/gcd2^,* and *Atm^tm1Awb/tm1Awb^*. Data points represent the percentage of micronucleated cells from individual mice. Data were compared using a one-way ANOVA and Tukey’s multiple comparisons test. (G) Representative images of nuclear shapes and corresponding solidity values (s) observed in *KIF18A* KO hTERT-RPE1 cells. (H) Box and whisker plot of nuclear solidity values measured in control (n = 553) and *KIF18A* KO (n= 634) cells. Data distributions were compared using a Kolmogorov-Smirnov t-test. (I) Plot of percentage of nuclei with solidity values two standard deviations below the average in control cells. Data were compared using a Chi-square test. In all panels, asterisks (*) indicate p<0.01.

In addition to micronuclei, KIF18A-deficient cells contain abnormally shaped primary nuclei with a lobed appearance (Figure 3G). To quantify this phenotype, we measured the solidity of DAPI-stained nuclei in control and *KIF18A*-KO hTERT-RPE1 cells. *KIF18A*-KO cells displayed a significant increase in the percentage of primary nuclei with solidity values that differed by more than two standard deviations from the mean solidity of control cells (Figure 3 G-I). Thus, loss of KIF18A activity disrupts the normal convex shape of primary nuclei in hTERT-RPE1 cells.

### Micronuclei and abnormal nuclear shapes form as KIF18A-deficient cells exit mitosis

To determine if the interphase nuclear organization abnormalities observed in KIF18Adeficient cells result from mitotic defects, we analyzed when micronuclei and lobed primary nuclei form relative to cell division in live *KIF18A*-KO cells expressing H2B-GFP (Figure 4, Figure S3, and Supplemental Movies S1-S3). We found that 9.8% of daughter cells formed micronuclei as chromatin decondensation occurred at the end of mitosis in *KIF18A*-KO cells compared to 1.2% in hTERT-RPE1 control cells (Figure 4 A-B and Table S2). This percentage is comparable to the fraction of cells with micronuclei observed in fixed *KIF18A* KO cells (Figure 3C). While the shape of primary nuclei in cells entering mitosis (“mother cells”) was comparable between *KIF18A*-KO and control cells (Figure 4 C-D), *KIF18A*-KO daughter cells formed a significantly higher fraction of abnormal nuclei than control cells during mitotic exit (Figure 4 E-F). The percentage of daughters with abnormal nuclear shapes is strikingly similar to the percentage of cells with abnormal nuclear shapes observed in fixed samples (Figure 3 H-I). These data strongly suggest that mitotic chromosome alignment defects underlie the majority of the nuclear organization abnormalities observed in interphase cells lacking KIF18A function.

**Figure 4.**
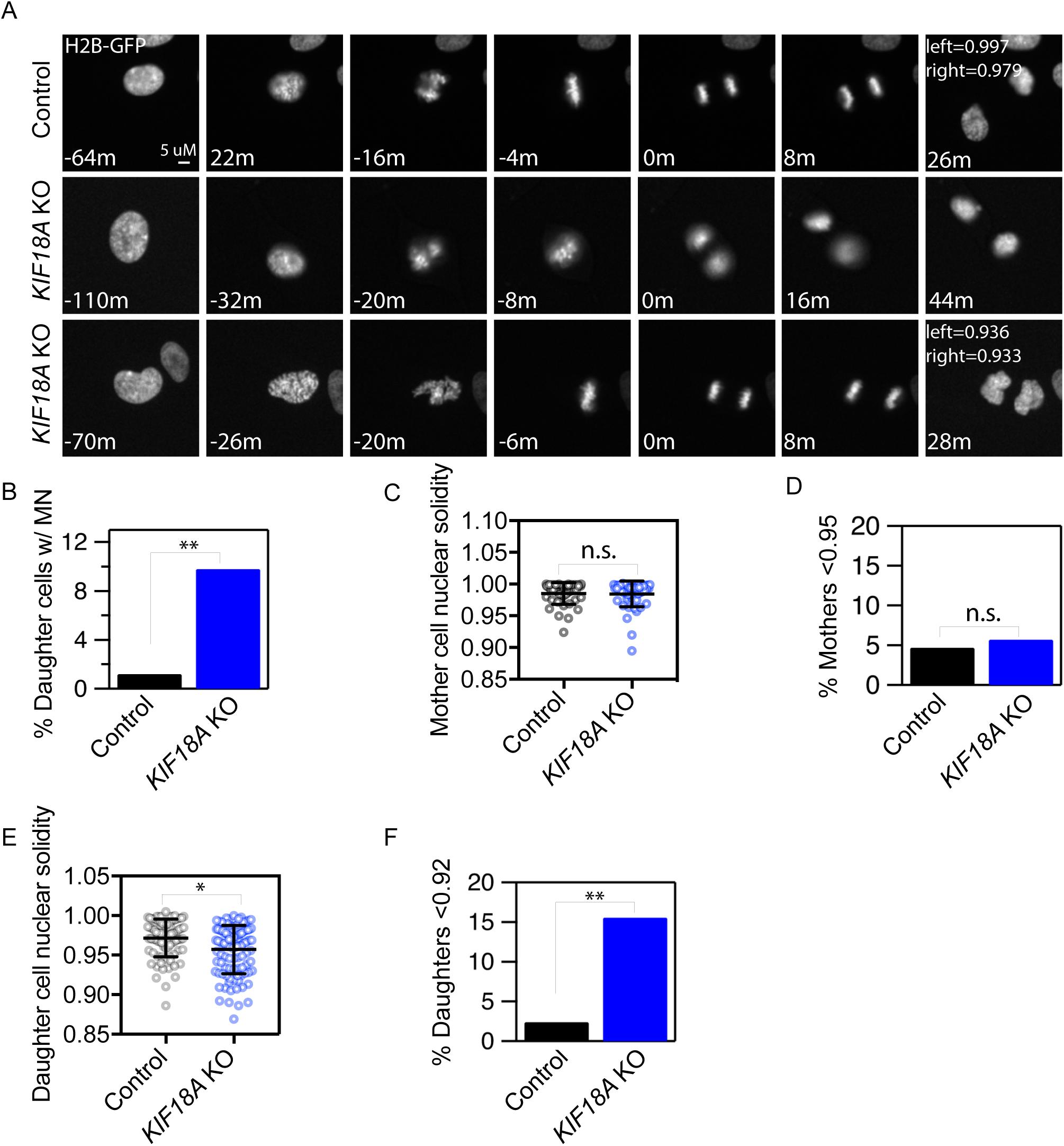
Micronuclei and abnormal nuclear shapes form as KIF18A-deficient cells exit mitosis. (A) Representative stills from time-lapse images of control and *KIF18A* KO hTERTRPE1 cells expressing histone H2B-GFP. Note that micronuclei (middle panels) and lobed primary nuclei (bottom panels) form as *KIF18A* KO cells exit mitosis. (B) Plot of the percentage of daughter cells that form micronuclei during mitotic exit. Data were compared via Chi-square test, asterisks (**) indicate p<0.01 (C) Quantification of mother cell nuclear solidity measured 60 minutes prior to metaphase. Error bars indicate mean and standard deviation. Data were compared via Kolmogorov-Smirnov t-test, p>0.90. (D) Percentage of mother cell nuclear solidity values less than two standard deviations from the average control solidity. Data were compared via Chi-square test, p>0.90. (E) Quantification of daughter cell nuclear solidity 20 minutes after initial chromatin decondensation. Data were compared via Kolmogorov-Smirnov t-test, asterisks (*) indicate p<0.05. (F) Percentage of daughter cell nuclear solidity values two standard deviations below the average control solidity. Data were compared using Chi-square tests, asterisks (**) indicate p<0.01.

### Disruption of chromosome alignment leads to inter-chromosomal compaction defects and abnormal nuclear envelope reformation

To gain an understanding of how nuclear organization problems result from mitotic defects in *KIF18A*-deficient cells, we compared chromosome segregation and nuclear envelope reformation in the presence and absence of KIF18A function. These analyses revealed that anaphase chromosomes are more broadly distributed in both *KIF18A*-KO and *KIF18A*-KD cells compared to controls (Figure 5 A-C). Chromosome distributions during anaphase were quantified by measuring the standard deviation of centrosome-to-kinetochore distances within each half spindle. *KIF18A*-KD and *KIF18A*-KO hTERT-RPE1 cells display a greater variance in pole-to-kinetochore distances than control cells (Figure 5 B-C). These data indicate that *KIF18A* is required for inter-chromosomal compaction during anaphase.

**Figure 5.**
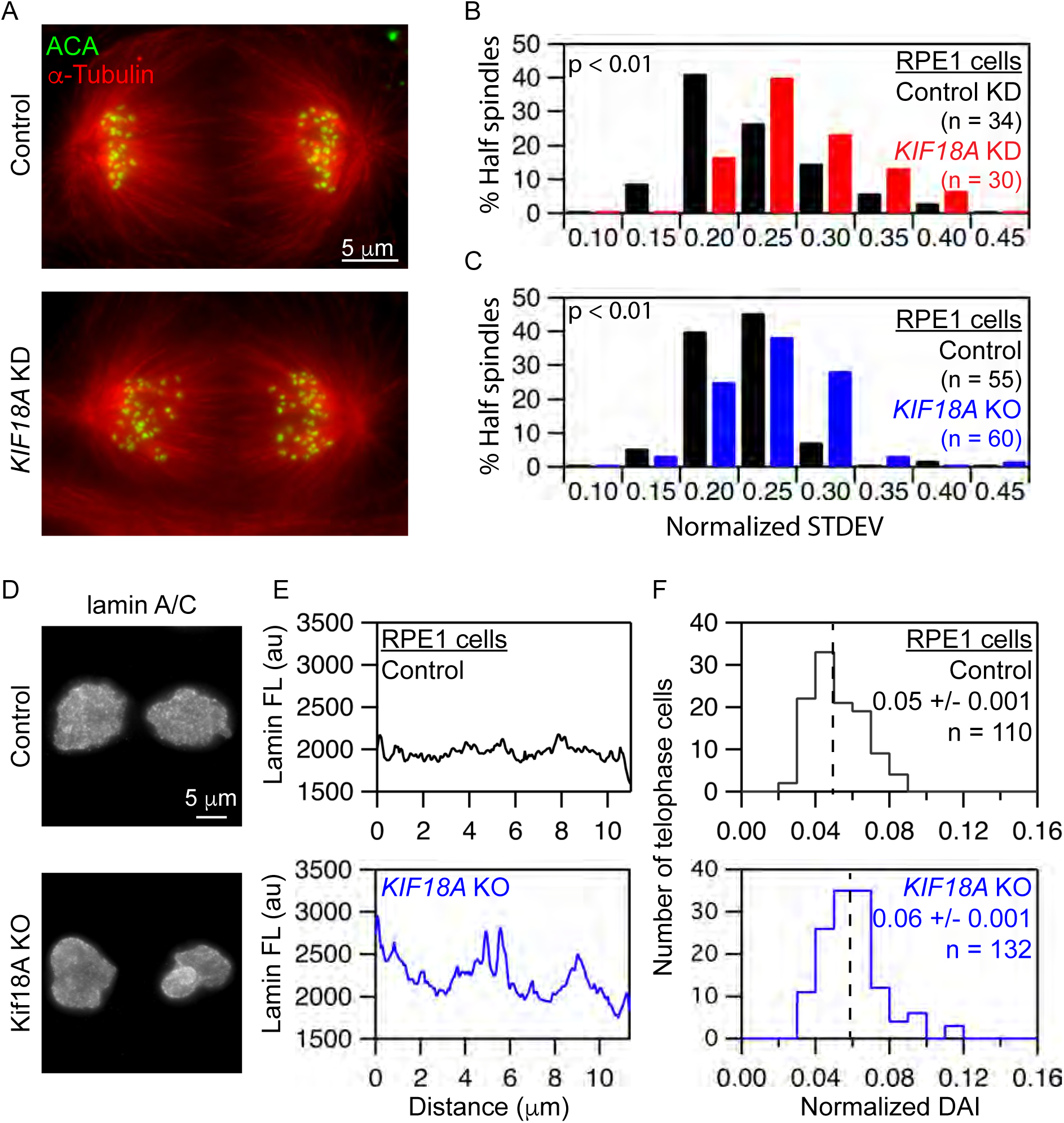
Loss of KIF18A function and chromosome alignment disrupts interchromosomal compaction during anaphase and lamin A/C distribution during telophase. (A) Representative images of anaphase cells fixed and stained for α-tubulin (red) and centromeres (ACA, green). (B-C) Histograms of centromere to pole distance variance (calculated as standard deviation) among all centromeres within a half spindle of (B) control and *KIF18A* siRNA treated hTERTRPE1 cells or (C) control and *KIF18A* KO hTERT-RPE1 cells. Data were compared using a Kolmogorov-Smirnov t-test, p<0.01. (D) Representative images of control and *KIF18A* KO telophase cells labeled with Lamin A/C antibodies. (E) Plots of Lamin A/C fluorescence profiles along the long axis of telophase nuclei from control and *KIF18A* KO hTERT-RPE1 cells. (F) Histograms of Lamin A/C fluorescence variance in telophase nuclei calculated as the deviation from the average intensity (DAI) in control and KIF18A KO cells. Distributions of DAI were compared using a Kolmogorov-Smirnov t-test, p<0.01.

Defects in inter-chromosomal compaction are predicted to disrupt proper nuclear envelope reformation during telophase (Mora-Bermúdez et al., 2007). To test this, we assayed the distribution of lamins in fixed telophase cells by measuring the deviation of average fluorescence intensity (DAI) along the long axis of each DNA mass in cells immunofluorescently labeled for lamin A/C (Figure 5 D-F). *KIF18A*-KO cells showed a significantly increased variance in lamin A/C signal along the chromosome mass compared to controls, indicating that lamin is abnormally distributed during nuclear envelope reformation in the absence of KIF18A.

### Lagging chromosomes in KIF18A-depleted cells travel longer distances in anaphase

To determine the basis of inter-chromosomal compaction and nuclear organization defects in KIF18A-deficient cells, we imaged chromosome and kinetochore dynamics during mitosis with high temporal resolution. Analyses of H2B-GFP expressing anaphase cells confirmed inter-chromosomal compaction defects during anaphase following *KIF18A*-KD (Figure 6A). Of 22 cells imaged from metaphase to telophase, 18 displayed obvious anaphase compaction defects and lagging chromosomes (82%). In 11.4% of the daughter cells resulting from *Kif18A*-KD cell divisions (5 of 44), we observed that a lagging chromosome was excluded from the primary nucleus and formed a micronucleus (Figure 6A). These data suggest that, in the absence of KIF18A function, micronuclei form around lagging mitotic chromosomes that arrive late to the poles during anaphase.

**Figure 6.**
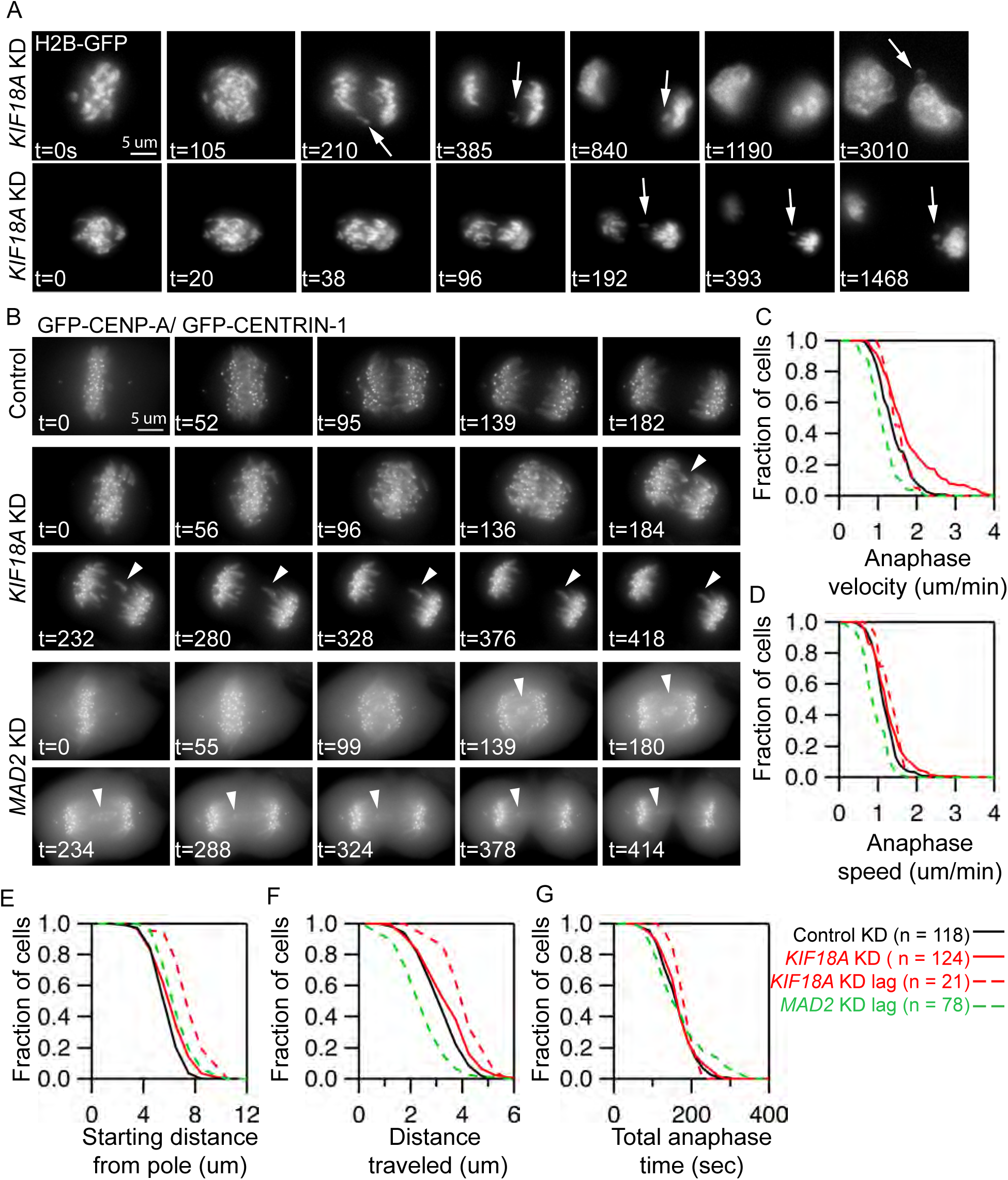
In the absence of chromosome alignment, micronuclei form around lagging chromosomes that travel long distances during anaphase. (A) Stills from time-lapse imaging of *KIF18A*-depleted cells expressing histone H2B-GFP. Arrows indicate lagging chromosomes that are excluded from the primary DNA mass and form micronuclei. (B) Representative images of live cells stably expressing GFP-CENP-A and GFP-CENTRIN-1 treated with control, *KIF18A*, or *MAD2* siRNAs. Arrowheads indicate lagging chromosomes, (C-G) Survival plots of (C) poleward anaphase velocity (um/min), (D) poleward anaphase speed, (E) starting distance from the pole, (F) distance traveled, and (G) total anaphase time for kinetochores in each experimental condition indicated. Dashed lines indicate the behavior of lagging chromosomes in *KIF18A* and *MAD2* siRNA treated cells. Data were compared using a Kruskal-Wallis test with post-hoc Dunn’s multiple comparison tests. The starting distance from the pole and distance traveled for lagging chromosomes in *KIF18A* siRNA cells were significantly different than those in control siRNA cells or total kinetochores in KIF18A KD cells (p < 0.01). The anaphase velocity, speed, and distance traveled for lagging chromosomes in *MAD2* KD cells are significantly different than those of kinetochores in control siRNA cells (p < 0.01).

To understand the underlying defect causing lagging chromosomes in KIF18A-deficient cells, we tracked the movements of fluorescently labeled kinetochores in *KIF18A* and control siRNA treated cells stably expressing EGFP-CENP-A and EGFP-Centrin-1 (Figure 6B) (Magidson et al., 2011). These studies revealed that lagging chromosomes in *KIF18A*-KD cells moved to the poles with normal speeds and velocities but began anaphase at significantly increased distances from the pole, and therefore, moved longer distances during anaphase (Figure 6 C-F). However, the time that chromosomes were moving to the poles during anaphase was not changed in *KIF18A*-KD cells relative to controls. In contrast, we found that lagging kinetochores in *MAD2*-KD cells move significantly slower but travel similar distances to kinetochores in control cells (Figure 6 C-F). The slow movements of lagging chromosomes in *MAD2*-KD cells are likely due to merotelic attachments that are not corrected before the chromosomes prematurely segregate (Cimini et al., 2003). The differences between the movements of lagging chromosomes in *KIF18A*-KD and *MAD2*-KD cells suggest that merotelic attachments do not significantly contribute to the defects in *KIF18A*-KD cells. This conclusion is further supported by the low percentage of late anaphase *KIF18A*-KD cells with midzone lagging chromosomes (2.4%, n = 286 cells), which is comparable to the number observed in late anaphase control cells (1.0%, n = 297 cells, p = 0.18). Late anaphase lagging chromosomes result from severe merotelic attachments (Cimini et al., 2004). Taken together, our data suggest that chromosome alignment defects in *KIF18A*-KD cells abnormally position chromosomes at anaphase onset, in turn leading to inter-chromosomal compaction defects and lagging chromosomes during segregation.

### A p53-dependent checkpoint reduces division of micronucleated KIF18A-deficient cells

Micronuclei have been identified as markers of chromosome instability in tumors and their presence can lead to chromothripsis, a phenomenon involving extensive structural rearrangements within a single chromosome (Crasta et al., 2012; Zhang et al., 2015). Interestingly, *Kif18a* mutant mice form micronuclei *in vivo* (Figure 3F) but have been reported to be tumor-resistant rather than predisposed to tumor formation (Nagahara et al., 2011; Zhang et al., 2010; Zhu et al., 2013). This raises the question of whether KIF18A-deficient cells with micronuclei are able to continue to proliferate. We used long-term live cell imaging to measure the rate of cell division for micronucleated cells in Kif18A-KO and control hTERTRPE1 populations. The percentage of micronucleated cell divisions was calculated as a function of the fraction of micronucleated cells in the population. These analyses revealed that micronucleated hTERT-RPE1 control and *KIF18A*-KO cells display reduced division rates of 37.5% and 32.9%, respectively (Figure 7 A-B and Table S2). These division rates are consistent with those measured following induction of micronuclei and chromosome segregation errors via treatments that do not cause mitotic arrest (Soto et al., 2017). Previous studies have indicated that micronucleus formation can lead to cell cycle arrest via a p53- dependent checkpoint (Sablina et al., 1998; Thompson and Compton, 2010). We found that the rate of division for micronucleated control and *KIF18A*-KO cells increases 2 to 3-fold when p53 is depleted by siRNA treatment (Figure 7B, Table S2). These data indicate that micronuclei limit proliferation in *KIF18A*-KO cells, at least in part, through a p53-dependent mechanism.

**Figure 7.**
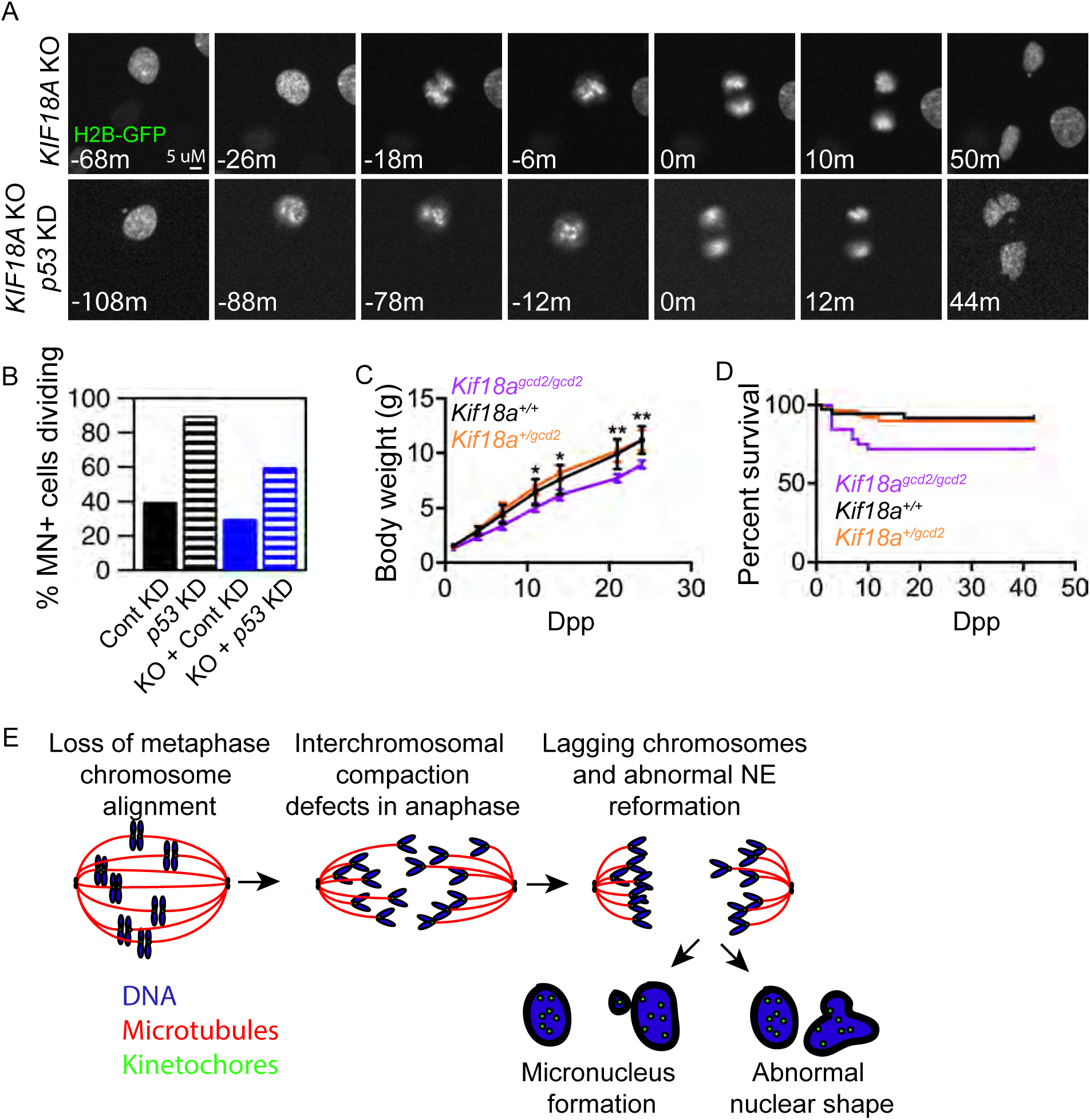
A p53-dependent mechanism limits the division of micronucleated *KIF18A* KO cells. (A) Still frames from time-lapse analyses of dividing histone 2B-GFP (H2B-GFP) expressing *KIF18A* KO cells and *KIF18A* KO cells treated with *p53* siRNAs. (B) Plot of the percent of micronucleated cells that enter mitosis in control, *p53* KD, *KIF18A* KO, or *KIF18A* KO + *p53* KD hTERT-RPE1 cells. (C) Plot of body weights measured at the indicated days postpartum (Dpp) for each genotype listed. (D) Survival plot for mice of the indicated genotypes as a function of Dpp. (E) Model for abnormal nuclear formation in the absence of chromosome alignment (see Discussion text for details).

### *Kif18A* mutant mice display reduced growth rates and increased pre-wean mortality

Consistent with the reduction in proliferation observed in *KIF18A*-KO hTERT-RPE1 cells, we previously found that MEFs derived from *Kif18A^gcd2/gcd2^* mice grow slower than those from wild type or *Kif18A^gcd2/+^* mice, despite progressing through mitosis with normal timing (Czechanski et al., 2015). Furthermore, the percentage of homozygous mutant mice found in F2 litters (18%) was slightly below the expected Mendelian ratio, indicating lethality prior to genotyping at wean (Czechanski et al., 2015). To determine if *Kif18A^gcd2/gcd2^* mice display growth defects, body weight was measured over a time course of 24 days post-partum (dpp). The body weights and growth rate of *Kif18A^gcd2/gcd2^* mice were significantly reduced compared to either wild type or *Kif18A^gcd2/+^* littermates (Figure 7C). *Kif18A^gcd2/gcd2^* mice also exhibited increased mortality during this time period, as predicted by our previous observations (Figure 7D). These phenotypes suggest that proper chromosome alignment is required for normal growth and viability during post-natal development.

## DISCUSSION

Chromosome compaction during anaphase is required to limit the frequency of lagging chromosomes and to promote formation of a single nucleus around all chromosomes at the end of cell division (Mora-Bermúdez et al., 2007; Ohsugi et al., 2008). This involves both the clustering of chromosomes together (interchromosomal compaction) and axial shortening of individual chromosome arms (intrachromosomal compaction). While mechanisms contributing to intrachromosomal compaction have been reported (Mora-Bermúdez et al., 2007; Ohsugi et al., 2008), the molecular control of interchromosomal compaction is less understood. Our studies of chromosome alignment-deficient cells, which lack KIF18A, indicate that a key function of metaphase chromosome alignment is to provide this clustering prior to anaphase onset, setting the stage for normal nuclear envelope assembly. In this model, a primary function of chromosome alignment is to equalize the distances that chromosomes travel during anaphase segregation. This, in turn, organizes chromosomes into a compact mass during anaphase and promotes uniform templating of nuclear envelope components during telophase, resulting in the formation of a single, ovular nucleus in each daughter cell at the completion of division.

We observed that the vast majority of chromosome-alignment defective cells displayed lagging chromosomes and interchromosomal compaction defects, yet a much smaller fraction of daughter cells formed micronuclei or abnormally shaped primary nuclei. The presence of mechanisms that spatially coordinate nuclear envelope formation with chromosome segregation can explain this discrepancy. For example, an Aurora B kinase gradient at the midzone of anaphase cells inhibits nuclear envelope assembly until chromosomes have segregated to the poles (Afonso et al., 2014). This mechanism allows most lagging chromosomes to integrate into the primary nucleus, reducing the frequency of micronucleus formation. However, lagging chromosomes can still be excluded from the main nucleus even in the presence of this surveillance, accounting for the low but consistent percentage of micronucleated cells formed in the absence of chromosome alignment (Afonso et al., 2014).

Chromosome alignment per se does not appear to be required for proper kinetochore microtubule attachments in normal mammalian somatic cells. Despite severe disruption of chromosome alignment, KIF18A-deficient somatic cells from mouse and human progress through mitosis with relatively normal timing compared to the long mitotic delays observed in germ cells and HeLa cells lacking KIF18A function. These data suggest that chromosome alignment-defective MEFs and hTERT-RPE1 cells establish attachments that satisfy the spindle assembly checkpoint with kinetics similar to or slightly slower than control cells, respectively. Furthermore, the lagging chromosomes resulting from chromosome alignment defects displayed distinct behaviors compared to those that result from merotelic attachments, which typically segregate with reduced velocities and often stall at the midzone during anaphase (Cimini et al., 2004). These data strongly support the conclusion that chromosome alignment defects, rather than abnormal attachments, lead to nuclear formation problems.

Our analyses of chromosome copy number also suggest that metaphase alignment may not be necessary for the equal segregation of chromosomes. However, it is surprising that a population of cells where ~10% are micronucleated display no change in chromosome copy number. We believe this apparent discrepancy can be explained by sensitivity limitations of standard assays for chromosome copy number. If it is assumed that there is no bias in selecting which chromosomes form micronuclei, any particular chromosome would be micronucleated in ~0.4% of KIF18A deficient cells (10% of cells with MN/ 23 chromosomes). This level of copy number change would not be detectable in our FISH analyses of primary nuclei. Furthermore, because micronucleated KIF18A-deficient cells display a low rate of division, any copy number defects in these cells are unlikely to impact our analyses of mitotic chromosome spreads. Thus, we acknowledge that there may be a low level of aneuploidy induced by loss of KIF18A function and chromosome alignment that was not detected in our assays. However, our data indicate that chromosomes are equally segregated in the majority chromosome alignment defective cells, supporting the conclusion that alignment is largely dispensable for maintaining chromosome copy number.

We found that the cellular defects resulting from loss of chromosome alignment have longterm consequences that negatively impact cell proliferation, early mammalian growth, and survival. At least a portion of micronucleated cells resulting from chromosome alignment defects undergo a p53-dependent cell cycle arrest. This response is consistent with those observed in cells containing micronuclei formed as a result of genomic instability (Sablina et al., 1998; Soto et al., 2017; Thompson and Compton, 2010) and can account for the reduced proliferation of *Kif18a^gcd2/gcd2^* MEFs and reduced growth of *Kif18a^gcd2/gcd2^* mice. Survival is also reduced in some *Kif18a^gcd2/gcd2^* mice, demonstrating that loss of chromosome alignment can compromise viability. The reduced fitness of *Kif18a* mutant mice in a highly controlled laboratory environment suggests that mice harboring defects in chromosome alignment may be sensitized to additional genetic or environmental factors that are detrimental to the mitotic / meiotic spindle, or to genome integrity. In fact, the intermediate phenotypes we observed in mice and in cell lines heterozygous for *Kif18a* mutations show that even subtle defects in chromosome alignment contribute to semidominant phentoypes. Therefore, it will be important to determine if *Kif18a* mutant mice are predisposed to aneuploidy and to explore long term consequences of this in the context of stem cell exhaustion, tumorigenesis, and overall population fitness.

In summary, our data indicate that the organization of chromosomes at the spindle equator during metaphase primarily functions to cluster chromosomes for segregation and ensure the proper formation of a single nucleus in each daughter cell. In the absence of this function, nuclear organization defects, such as micronuclei and abnormal nuclear shape, reduce postnatal mammalian growth and survival.

## EXPERIMENTAL PROCEDURES

### Animal ethics statement

All procedures involving mice were approved by The Jackson Laboratory’s Institutional Animal Care and Use Committee and performed in accordance with the National Institutes of Health guidelines for the care and use of animals in research. A cryorecovery of CAST. 129S1(B6)-*Kif18a^gcd2^*/Jcs (RRID:MMRRC_034325-JAX) (Czechanski et al., 2015) x C57BL/6J (RRID:IMSR_JAX:000664) was used to establish a breeding colony, which was maintained by sibling intercrossing.

### Cell Culture and Transfections

hTERT-RPE-1 cells (ATCC, Manassas, Virginia) were maintained at 37°C with 5% CO2 in MEM-α (Life Technologies, Carlsbad, CA) containing 10% fetal bovine serum (FBS; Life Technologies) and antibiotics. For time course experiments, cells were plated in a 6-well dish and transfected with 150pmol of siRNAs using RNAiMAX (Life Technologies) following the manufactures instructions. Fresh siRNAs were added every 48hr. Cells were then seeded on 25 mm coverslips (Electron Microscopy Sciences, Hatfield, PA) for imaging. For other fixed cell assays, cells were seeded on 12mm acid-washed coverslips and transfected with 30 pmol siRNA complexed with RNAiMAX. For live cell imaging, cells were seeded in 35 mm poly-L-lysine coated glass bottom dishes (MatTek, Ashland, MA) 24 hr before addition of siRNA. Plasmid transfections were carried out using a Nucleofector 4D system (Lonza, Walkersville, MD).

### Mouse embryonic fibroblast derivation and culture

For derivation of mouse embryonic fibroblasts, F3 embryos were harvested and euthanized at 12.5-14.5 days post coitum (dpc, copulation plug = 0.5 dpc). Tissue samples were collected for genotyping and each embryo was processed individually as previously described to avoid cross contamination between genotypes (Czechanski et al., 2015; 2014). Individual, eviscerated, decapitated embryos were washed in cold PBS, transferred to clean 100 mm dishes, macerated with forceps in 3-5 ml of 0.05% Trypsin/EDTA (ThermoFisher, cat# 25300120), and then dissociated through 18G needles. Tissue suspensions were transferred to 15 ml conical tubes and an equal volume of pre-warmed 37C, MEF medium (DMEM [ThermoFisher, cat#11960044], 10% fetal bovine serum [Lonza, cat#14-501 F, lot#0000217266], 100 U/ml penicillin / streptomycin [ThermoFisher, penicillin-streptomycin 1000U/ml, cat#15140122], 1X Glutamax [ThermoFisher, cat# 35050061], 0.2 µM filtered) was added. Large fragments were allowed to settle and cell suspensions were transferred to clean 15 ml conical tubes and centrifuged for 5 min. at 200 x g at room temperature. Cell pellets were resuspended in 5 ml of pre-warmed, 37C MEF media and plated onto clean 60mm tissue culture dishes (P0). Cultures were incubated at 37C, 5% CO_2_ for 2-4 days until 80% confluent. Cultures were then harvested in 0.05% trypsin/EDTA, aliquots were removed for genotype confirmation, and cultures were resuspended in freeze medium (10% dimethyl sulfoxide [DMSO]/10% FBS/80%MEF) at 0.5-1 × 10^6 cells / ml, dispensed into 1 ml aliquots in cryovials and then gradually frozen (−1°C/min) at −80C overnight in CoolCell freezing containers. Frozen vials were then immersed in liquid nitrogen for storage. Tissue and cell samples were genotyped by The Jackson Laboratory Transgenic Genotyping Service using an endpoint assay designed to detect the R308K missense “*gcd2*” mutation at Chr2: 109,908,059 G/T, GRCm38, mm10. Primer and oligo sequences are listed in the Key Resources Table.

### CRISPR targeting of Kif18A

A sgRNA guide sequence was designed against the 15^th^ exon in the C-terminus of Kif18A with the sequence 5’- CTAATGCCATCTCCCTTGAA (AGG) -3’. The PAM sequence is in parenthesis. Primers were designed with this sequence and cloned into pCR-Blunt with gBLOCK through site-directed mutagenesis, and confirmed by T7 sequencing.

For transfection of the complex, approximately 2 × 10^6^ RPE1 cells plated into a 60 mm dish with 3 mL MEM-α media (Thermo Fisher), supplemented with 10% FBS and 1% Penicillin/Streptomycin (Thermo Fisher) were treated with 1 µg pCR-Blunt sgRNA and 1 µg pX458 Cas9-GFP with 16 µL Lipofectamine LTX (Thermo Fisher), incubated for 20 minutes at room temperature in 250 µL OptiMeM (Thermo Fisher). Similar amounts of RPE1 cells were also transfected as negative and positive controls for FACS, respectively: 2 µg pCRBlunt sgRNA (neg) and 2 µg pMAX GFP (Lonza) each with 16 µL Lipofectamine LTX incubated for 20 minutes at room temperature in 250 µL OptiMeM. The treated RPE1 cells were incubated at 37°C for 48 hours.

The Cas9 + sgRNA treated cells were then strained and sorted for positive GFP signal using a BD FACSAria cell sorter equipped with 488 nm Coherent Sapphire laser. Positive and negative controls were used to place bins for determining the cell population desired for sorting. 1 µg/mL DAPI was also included in the sample as a live/dead marker. Cells with high GFP signal and low DAPI signal were selected and automatically placed 1 cell per well into 96 well plates containing 100 µL MEM-α media supplemented with 20% FBS. Cell colonies that grew were subsequently tracked and re-plated into larger well sizes periodically over a period of approximately 3 weeks. From a 24-well size dish, a colony was divided equally into 2 6-well sized wells. One well was used to continue growing the cell population, the other at confluency was lysed and the genome extracted using a Blood Mini Kit (Qiagen) following the Appendix B protocol for cultured cells.

Cell genomes were screened by PCR amplification of the exon region of interest using genomically specific primers designed with the BLAT bioinformatics design tool (UCSC Genome Bioinformatics). Forward primer: 5’- GTAATAAACTGGTCACTGACACCCAAACCC - 3’; Reverse primer: 5’- GGGTAATTTACACTTCGAGCTCTTGATGTCTTC -3’. The guide was designed with a unique restriction enzyme (BslI, New England Biolabs) cut site overlapping the Cas9 cleavage site, such that mutations to the genome would negate BslI cutting. PRC amplified genomes that did not cut were then sequenced using the forward primer above. The Kif18A deficient CRISPR line used in this study has frames shifts in each chromosome, resulting in premature truncations at amino acid residues 675 and 673.

### Plasmids and siRNAs

H2B-GFP was a gift from Geoff Wahl (Addgene plasmid # 11680). Cells were transfected with pools of siRNAs targeting the Kif18A sequences: 5’-UCUCGAUUCUGGAACAAGCag-3’ and 5’-CCACUUUAUGAAAUCCAGCtg-3’; the Mad2 sequences 5’- UAUUUUCCUCAUGUCAUCCtt-3’ and 5’-AGAUGGAUAUAUGCCACGCtt-3’; or scrambled negative control Silencer siRNA #1 (Life Technologies).

### Generation of anti-Kif18A Motor Domain antibodies

To generate antibodies against the motor domain of Kif18A, nucleotides 1-1089 of the coding region (GenBank Accession No. BC048347) were PCR amplified, and inserted into the BamHI/EcoRI sites of pGEX6P-1 (GE Healthcare). GST-Kif18A-NT was expressed in and purified from BL-21(DE3) cells using glutathione agarose (Sigma-Aldrich), and used to immunize rabbits (Cocalico). Kif18A-specific antibodies were affinity purified by passing anti-GST-depleted serum over Affi-Gel 10 (Bio-Rad, Hercules, CA) covalently coupled to GST-Kif18A-NT. Affinity-purified antibodies were dialyzed into phosphate-buffered saline (PBS), frozen in liquid nitrogen, and stored at −80°C.

### Cell Fixation and Immunofluorescence

hTERT-RPE1 cells were fixed in −20°C methanol (Fisher Scientific, Hampton, NH), 1% paraformaldehyde (Electron Microscopy Sciences) in −20°C Methanol, 4% Paraformaldehyde in 1X TBS [TBS; 2.6 mM Potassium Chloride, 24.7 mM Tris Base, and 136.8 mM NaCl at pH 7.4], or 0.5% Glutaraldehyde (Electron Microscopy Sciences) in Microtubule Stabilization Buffer [MTBS; 1X BRB80, 4mM EGTA, 0.5% Triton-X]. Cells were then washed in 1X TBS and blocked in antibody dilution buffer (Abdil) [Abdil: tris buffered saline pH7.4, 1% bovine serum albumin, 0.1% triton-X, 0.1% sodium azide] containing 20% goat serum. Cells were incubated with the following primary antibodies for 1 hour at room temperature in Abdil: mouse anti-α tubulin at 1ug/ml (sigma-aldrich, St. Louis, MO), mouse anti-γ-tubulin at 1 ug/ml (sigma-aldrich), rat anti-YL at 2 ug/ml (Millipore, Jaffrey, NH), mouse anti-human Lamin A/C at 1 ug/ml (Millipore), rabbit anti-cleaved caspase-3 at 1:100 (Cell Signaling, Danvers, MA), mouse anti-p53BP at 0.8ug/mL (Santa Cruz Biotechnology, Dallas, TX), or rabbit anti-GFP at 8ug/ml (Molecular Probes by Life Technologies, Eugene, OR). Cells were incubated with the following antibodies overnight at 4°C: human anti-centromere protein (ACA) (Antibodies Incorporated, David, CA) 2 ug/mL, rabbit anti-Kif18A (C-terminal) at 2 ug/mL (Bethyl Antibodies, Burlington, ON, Canada), and rabbit anti-Kif18A (N-terminal) at 3ug/ml. Cells were incubated for 1 hr at room temperature with goat secondary antibodies against mouse, rabbit, or human IgG conjugated to Alex Fluor 488, 594, or 647 (Molecular Probes by Life Technologies). Coverslips were mounted on glass slides with Prolong Gold antifade reagent plus 4′,6-diamidino-2-phenylindole (DAPI) (Molecular Probes by Life Technologies).

### Fluorescent In Situ Hybridization and Chromosome Spreads

For fluorescent in situ hybridization and chromosome spreads experiments, hTERT-RPE1 cells and mouse embryonic fibroblasts (MEFs) were dissociated and placed directly in hypotonic solution (0.8% sodium citrate and 0.4% potassium chloride) for 20 minutes, prefixed in Carnoy’s fixative (3:1 Methanol: Glacial Acetic Acid), pelleted, resuspended in Carnoy’s fixative, and washed four times with Carnoy’s fixative. For chromosome spreads, cells were dropped onto a glass slide inside a Thermotron Drying Chamber set at 25°C and 37% humidity for optimal spreading. Slides were dried and heated to 65°C for 30 minutes and either mounted with Prolong Gold containing DAPI, or trypsin banded and Giemsa stained. For fluorescent in situ hybridization, slides were immersed in 2X standard saline citrate (SSC) at room temperature for 2 minutes, then dehydrated in 70%, 85% and 100% ethanol for 1 minute each at room temperature and air dried. Commercial probes for locus specific regions 1p36, 1qter, 2 centromere, 3 centromere, 9q34, 22q11.2 (Cytocell) and for locus specific regions 7q11.2, 7q31, 8q21, 21q22, 14q32, 18q21(Abbott Molecular) were used. Probes were applied to samples, coverslipped, and rubber cemented. Slides and probes were co-denatured at 73°C for 2 minutes and hybridized at 37°C overnight in a hybridization chamber (Thermoscientific). Slides were then washed in 0.4X SSC/0.3%NP-40 for 2 minutes at 73°C, rinsed in 2X SSC/0.1% NP-40 for 1 minute at room temperature, counterstained with DAPI (Abbott Molecular) and coverslipped.

### Microscopy

Cells were imaged on a Nikon Ti-E inverted microscope (Nikon Instruments, Melville, NY) controlled by NIS Elements software (Nikon Instruments) with a Spectra-X light engine (Lumencore, Beaverton, OR), Clara cooled-CCD camera (Andor, South Windsor, CT), environmental chamber, and the following Nikon objectives: Plan Apo 20X DIC M N2 0.75NA, Plan Apo 40X DIC M N2 0.95NA, Plan Apo λ 60X 1.42NA, and APO 100X 1.49 NA.

### Live Cell Imaging

Cells were transferred into CO2-independent media with 10% FBS and 1% penicillin and streptomycin (Gibco by Life Technologies) for imaging via fluorescence or differential interference contrast (DIC) microscopy. For DIC imaging and long term imaging of EGFPH2B expressing cells, single focal plane images were acquired at 2-minute intervals with a 20X or 40X objective. For high temporal resolution imaging of chromosomes, optical sections were collected with 1.2 um spacing through the entire EGFP-H2B mass every 10s using a 60X objective. For imaging kinetochore movements in RPE-1 cells stably expressing EGFPCenp-A/EGFP-Centrin-1 (graciously gifted by Alexey Khodjakov) optical sections were collected at 1 um spacing every 5 seconds using a 60X objective (Magidson et al., 2011).

### Live Kinetochore Tracking

Maximum intensity projections of 3.6-6microns were made from optical sections of siRNA treated RPE-1 cells stably expressing EGFP-Cenp-A/EGFP-Centrin-1. Images were analyzed using MtrackJ in ImageJ. Centrin-1 was tracked throughout the movie from the end of metaphase through the end of anaphase B. Each kinetochores Cenp-A fluorescence position was tracked during pole ward movement. Kinetochore positions were normalized relative to the Centrin-1 track. Each kinetochore track was opened in Igor and individual kinetochore velocities were measure using the slope of the line from the beginning to end of anaphase. Speed is the velocity with pauses and reversals excluded. Pauses are a stop in pole ward kinetochore movement that occurred for 30 seconds or longer. Reversals are away from pole ward movement that occurred for 30 seconds or longer.

### Micronucleus Counts

For fixed cells, micronucleus counts were made using single focal plane images of DAPI stained cells. Image acquisition was started at a random site at the top edge of the coverslip and images were acquired every two fields of view with 40X objective. For live cell studies, the percentage of micronucleated cells was determined for each population by counting the number of cells with a pre-existing micronucleus in the first frame of each time-lapse data set. Micronuclei were defined as EGFP-H2B foci spatially separated from the primary nucleus, and migrating similarly alongside the primary nucleus of the cell. NIS Elements software (Nikon) was used to review the images and track micronucleus dynamics, as well as nuclear lobing. The percentage of cells dividing with a pre-existing micronucleus was calculated as the percentage of divisions wherein a cell with pre-existing micronucleus entered division, multiplied by the percentage of pre-existing micronuclei in that population (as estimated from the percentage of micronucleus-containing cells at the first frame of each field).

### Peripheral Blood Micronucleus Assay

Micronucleus assays of peripheral blood were conducted as previously described (Dertinger et al., 1996; Reinholdt et al., 2004). Peripheral blood was collected from the retro-orbital sinus of male and female laboratory mice, *Mus musculus* aged 12-18 weeks, N=5 for each Kif18a genotype, and N=4 for ATM. Strains used were CAST. 129S1(B6)-*Kif18a^gcd2^*/Jcs (RRID:MMRRC_034325-JAX) x C57BL/6J (RRID:IMSR_JAX:000664) F2 and F3 homozygous mutant and littermate wild type controls, as well as 129S6/SvEvTac-*Atm^tm1Awb^*/ J (RRID:IMSR_JAX:002753) (Barlow et al., 1996), and In addition to 75 µl of blood was immediately mixed with 100 µl of heparin and the mixture was then pipeted into 2 mls of ice-cold (−80C) 100% methanol with vigorous agitation to prevent clumping. Samples were stored at −80 C overnight before processing for flow cytometry. *Sample preparation and flow cytometry:* Each blood sample was removed from −80C storage and washed with 12 ml of autoclaved, ice-cold bicarbonate buffer (0.9% NaCl, 5.3 mM sodium bicarbonate, pH7.5), centrifuged at 500g for 5 min. and resuspended in a minimum of carryover buffer (~100 µl). 20 µl of each sample was added to a 5 ml polystyrene round-bottomed tube, and to each sample an 80 µl solution of CD71-FITC and RNAseA (1 mg/ml) was added. Additional control samples were CD71-FITC alone and an additional sample with bicarbonate buffer alone to which propidium iodide (PI) would be later added (see below). Cells were incubated at 4C for 45 minutes, washed with 2 ml cold bicarbonate buffer, and centrifuged as above. Cell pellets were stored on ice and then, immediately prior to flow cytometric analysis, resuspended in 1 ml of ice cold PI solution (1.25 mg/ml) to stain DNA. *Flow cytometry*: Samples were processed on a BD Bioscience LSRII fluorescence-activated cell sorter gated for FITC and PI, and set to collect 20,000 CD71 positive events at 5,000 events / sec. The CD71-FITC and PI control samples were used to calibrate for autofluorescence. Reticulocytes (Retic, CD71+, PI- [in the presence of RNAse A]), mature red blood cells (RBC, CD71-, PI-), micronucleated normochromatic erythrocytes (NCE-MN, CD71-, PI+) and micronucleated reticulocytes (Ret-MN, CD71+, PI+) were measured using FlowJo software. The total % of spontaneous micronuclei in NCE was NCE-MN/(NCE-MN + RBC)*100. Genotyping for *Kif18a^gcd2^* was as described above. Genotyping for *Atm^tm1Awb^* mice was performed by The Jackson Laboratory Transgenic Genotyping service using standard PCR.

### Anaphase Compaction Measurements

To measure anaphase compaction, optical sections were collected with 200 nm spacing through the entire spindle of fixed cells labeled with gamma-tubulin and ACA antibodies. The centroid of the gamma-tubulin focus in each half spindle was used as a reference point, and 3D distance measurements to each ACA-labeled kinetochore were made using ImageJ (NIH). The standard deviation of these distances was computed for each half spindle and used for comparison.

### Nuclear shape Analyses

For fixed cell analyses, single focal plane images DAPI-stained nuclei were acquired with a 20X objective. The shape of each nucleus was measured using the solidity measurement function in ImageJ (NIH). Nuclei were first thresholded and selected with the magic wand tool prior to shape measurements. For live cells expressing H2B-GFP, mother cell and daughter cell nuclear solidity were quantified from single focal plane images at the indicated time points relative to anaphase onset. Mother cell nuclear solidity measurements were made one hour prior to metaphase. Daughter cell nuclear solidity was measured twenty minutes after the second frame of chromosome de-condensation in telophase.

### Telophase Lamin Distribution Measurements

Single focal plane images of wild type and mutant Kif18A-/- RPE1 cells labeled with Lamin A/C were acquired through the central plane of the nucleus and opened in ImageJ. A line was drawn across the Lamin A/C signal along the long axis of the nucleus and the intensity of each pixel was imported into an Igor Pro (Wavemetrics, Portland, OR) macro. For each line the pixel intensity was normalized to max value from that scan. The normalized pixel intensity was then subtracted from average fluorescence for that line scan and the standard deviation of these values produced the deviation of the average intensity (DAI).

### Quantification of Chromosome Alignment

Chromosome alignment was quantified by calculating the full width at half maximum of ACA or DAPI fluorescence distribution along the pole-to-pole axis in ImageJ as described previously (Kim, 2014).

### Cleaved Caspase-3

Cleaved caspase-3 immunofluorescence was quantified using ImageJ (NIH). Cells expressing greater than two times the average fluorescence intensity measured in control cells were considered positive for cleaved caspase-3.

### Viability and growth, *Kif18a^gcd2^* mice

A cohort of 91 B6;CAST-*Kif18a^gcd2^* F3 mice representing all three *Kif18a^gcd2^* genotypes were monitored for growth. Animals were weighed every three days through wean and then monitored twice weekly through 8 weeks of age. Survival data were plotted using Prism 7 and survival curves were compared using Log-rank (Mantel-Cox) and Gehan-Breslow-Wilcoxon tests. Growth data were also plotted using Prism 7 and multiple t-tests at each time point were used to compare the mean weights across genotypes (Holm-Sidak, alpha=0.05)

## ACKNOWLEDGEMENTS

This work was supported by NIH grants GM121491 (to JS), R03 HD078485 (to LGR), and R01 GM086610 (to RO), by a Lake Champlain Cancer Research Organization grant (to JS), by Susan G Komen grant CCR16377648 (to JS), by a Leukemia and Lymphoma Career Development Award (to RO), and by a Vermont Space Grant Consortium fellowship (to LAS). The authors wish to thank Catherine Buck (University of Vermont Medical Center) for technical assistance with FISH and chromosome spread analyses.

## AUTHOR CONTRIBUTIONS

Conceptualization, LGR and JS; Methodology, CLF, HLHM, LAS, WM, CB, AC, MT, RO, LGR and JS; Investigation, CLF, HLHM, LAS, WM, CB, AC, DM, MT, LGR and JS; Resources, HLHM, WM, CB, AC, RO, LGR, and JS; Writing-Original Draft, CLF, HLHM, LAS, LGR, and JS; Writing-Review & Editing, CLF, HLHM, LAS, RO, LGR, and JS; Visualization, CLF, HLHM, LAS, LGR, and JS; Supervision, LGR and JS.

## SUPPLEMENTAL MOVIES

Movie S1. (related to Figure 4) Time-lapse movie of normal chromosome division in an hTERT-RPE1 cell expressing H2B-GFP.

Movie S2. (related to Figure 4) Time-lapse movie of micronucleus formation following division of a *KIF18A* KO hTERT-RPE1 cell expressing H2B-GFP.

Movie S3. (related to Figure 4) Time-lapse movie of abnormal nuclear shape formation following division of a *KIF18A* KO hTERT-RPE1 cell expressing H2B-GFP.

## FIGURE LEGENDS

**Figure S1.**
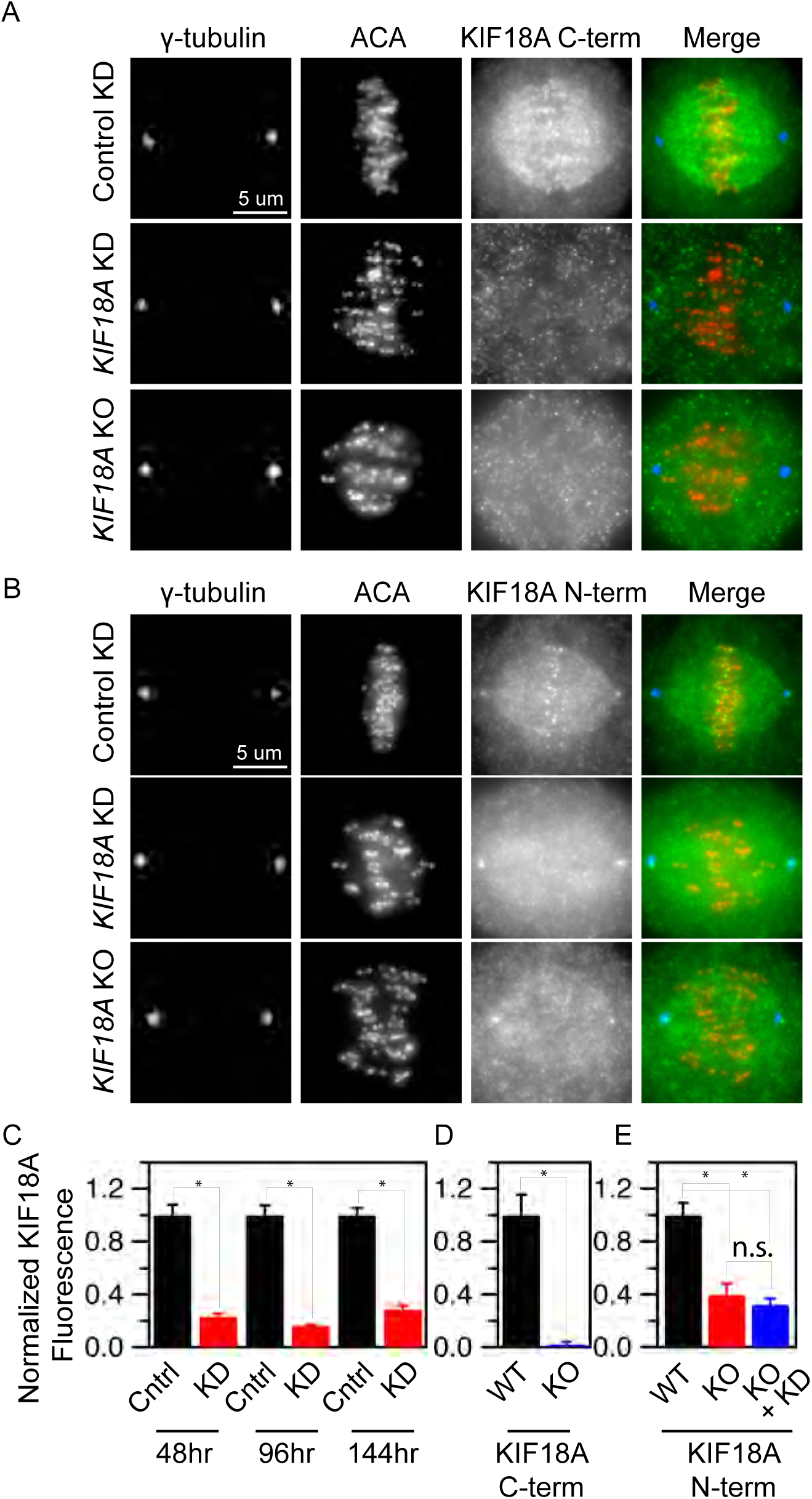
(related to Figure 1) *KIF18A* protein is undetectable on spindles in *KIF18A* KO hTERT-RPE1 cells. (A-B) Representative images of metaphase hTERT-RPE1 cells from the indicated conditions labeled for spindle poles (γ-tubulin), kinetochores (ACA), and KIF18A, using antibodies against either the KIF18A C-terminus (A) or N-terminus (B). Spindle pole staining from KIF18A N-term antibodies in non-specific. (C-D) Quantification of KIF18A immunofluorescence from *KIF18A* KD cells (C) and *KIF18A* KO cells (D-E). Note that treating *KIF18A* KO cells with *KIF18A* siRNAs (KO + KD) does not reduce KIF18A immunofluorescence levels.

**Figure S2.**
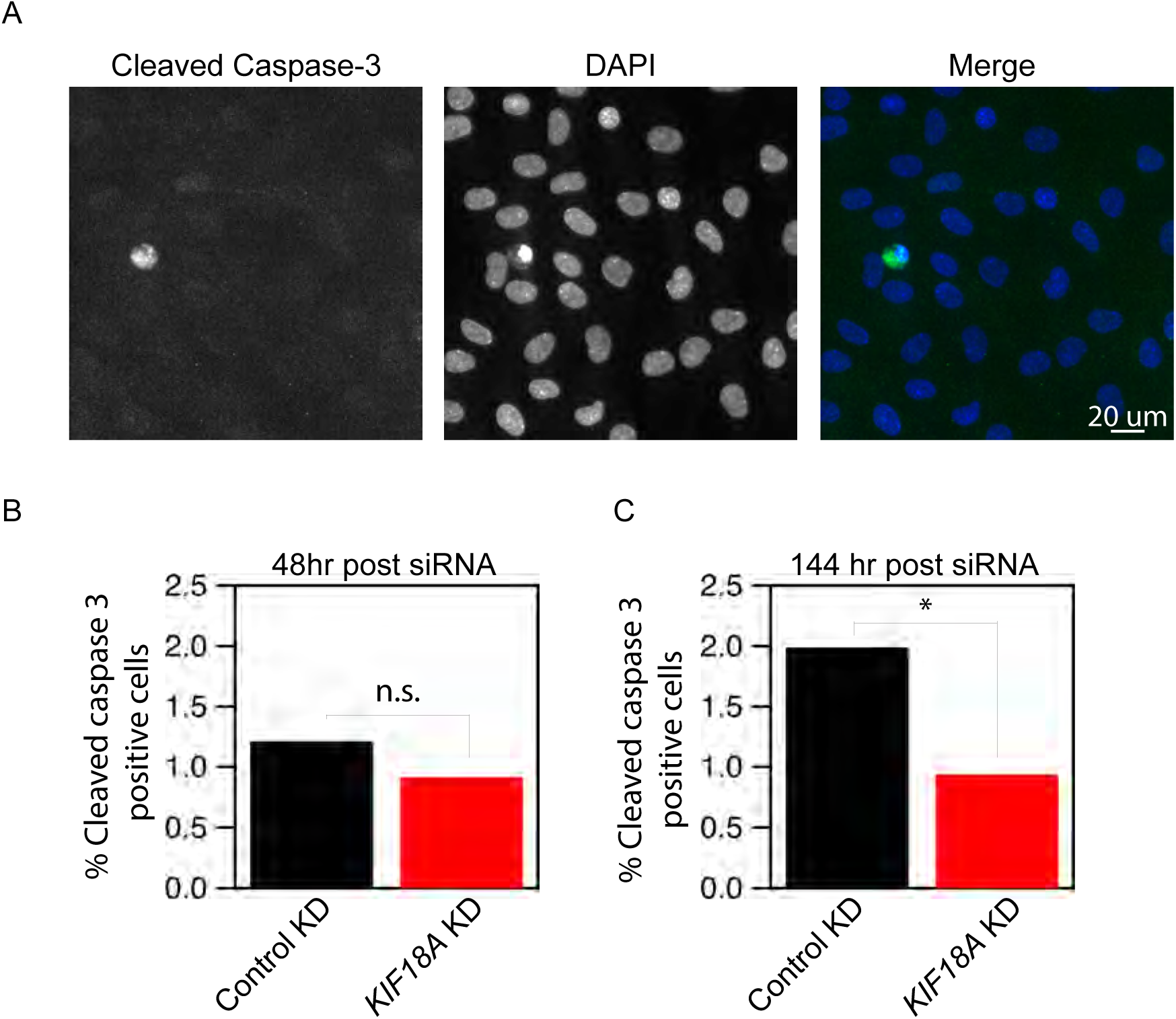
(related to Figure 2) Loss of KIF18A function does not lead to increased apoptosis. (A) Representative images of KI F 18A KD cells stained for the apoptotic marker Cleaved Caspase-3. (B-C) Plots indicating the percentage of cleaved caspase-3 positive cells in control and KIF18A KD cells (B) 48 hours and (C) 144 hours following siRNA treatment. Data were compared via Chi-square test. * indicates p < 0.05.

**Figure S3.**
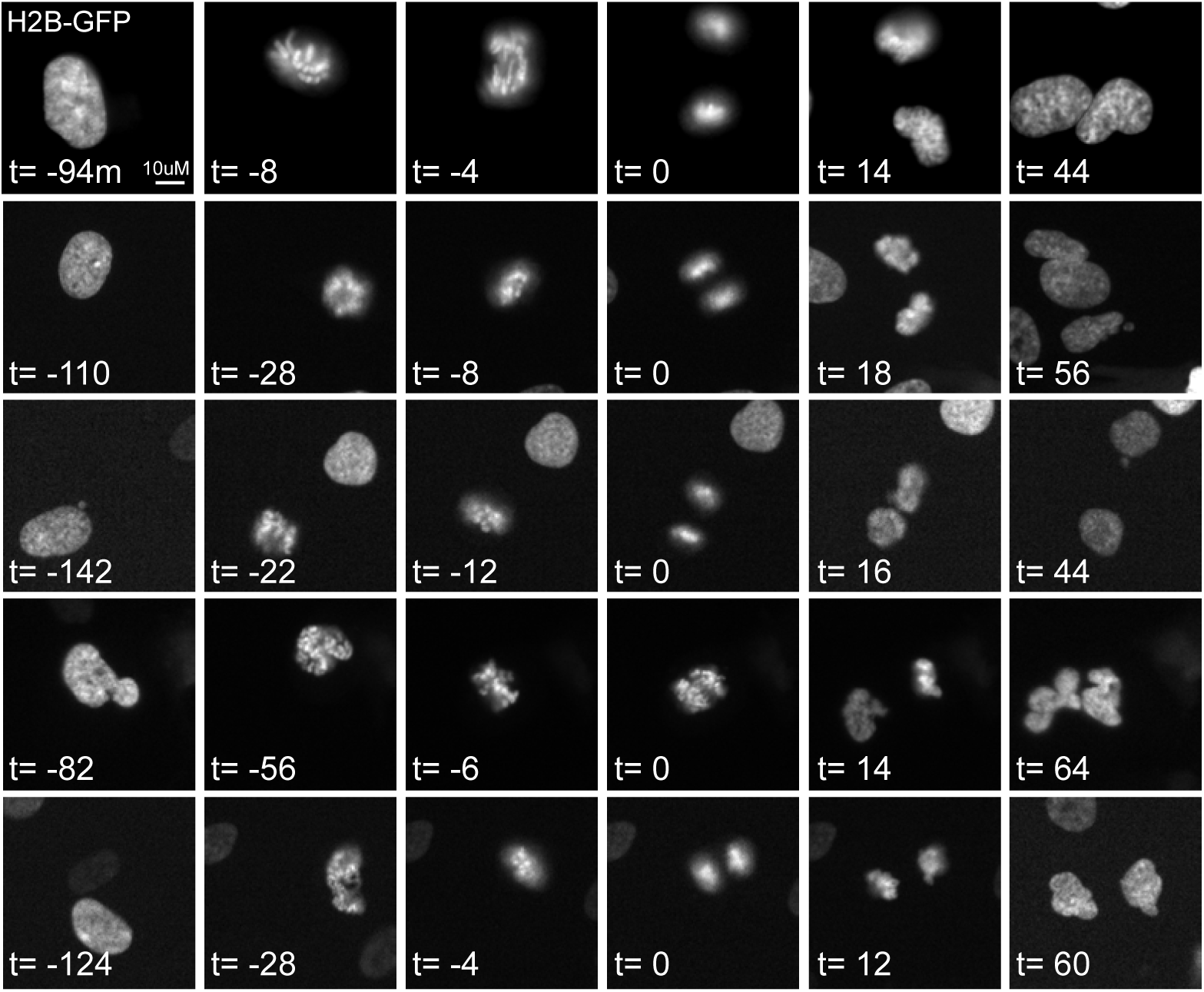
(related to Figure 4) Interphase nuclear defects occur as *KIF18A* KO cells exit mitosis. Panels display representative stills from time-lapse imaging of KIF18A KO cells transfected with histone-2B-GFP (H2B-GFP). Abnormal nuclear shapes and micronuclei are apparent within 1 hour of anaphase (time = 0).

**Table S1.** (related to Figure 2) Summary of fluorescent in situ hybridization data.

| **FISH probe** | **Control %**<br>**Aneuploid** | **Kif18A KD %**<br>**Aneuploid** | **<u>Chi-square p-value</u>** |
| --- | --- | --- | --- |
| **1qter** | 1.90(28/1472) | 1.97(29/1471) | 0.89 |
| **1p36** | 2.18(32/1468) | 2.46(36/1464) | 0.62 |
| **Centromere 3** | 3.59(52/1448) | 3.38(49/1451) | 0.76 |
| **Centromere 2** | 2.46(36/1464) | 2.95(43/1457) | 0.42 |
| **7q31** | 2.95(43/1457) | 4.02(58/1442) | 0.13 |
| **7q11.2** | 2.53(37/1463) | 3.02(44/1456) | 0.43 |
| **8q21.3** | 3.23(47/1453) | 2.25(33/1467) | 0.11 |
| **21q22.1** | 3.45(50/1450) | 3.45(50/1450) | 1.00 |
| **22q11.2** | 2.81(41/1459) | 2.67(39/1461) | 0.82 |
| **9.q34** | 3.66(53/1447) | 3.23(47/1453) | 0.54 |
| **14q32** | 1.35(20/1480) | 2.11(31/1469) | 0.12 |
| **18q21** | 1.69(25/1475) | 1.83(27/1473) | 0.78 |
|  | **Control %**<br>**Aneuploid** | **MAD2KD %**<br>**Aneuploid** | **<u>Chi-square p-value</u>** |
| **1qter** | 1.69(25/1475) | 6.61(93/1407) | <0.0001 |
| **1p36** | 1.83(27/1473) | 6.69(94/1406) | <0.0001 |
| **8q21.3** | 2.04(20/1470) | 3.95(57/1443) | 0.0033 |
| **21q22.1** | 1.91(20/1462) | 4.24(61/1439) | 0.0004 |
| **14q32** | 1.63(24/1476) | 3.02(44/1456) | 0.0142 |
| **18q21** | 2.88(42/1458) | 5.04(72/1428) | 0.0042 |

**Table S2.**
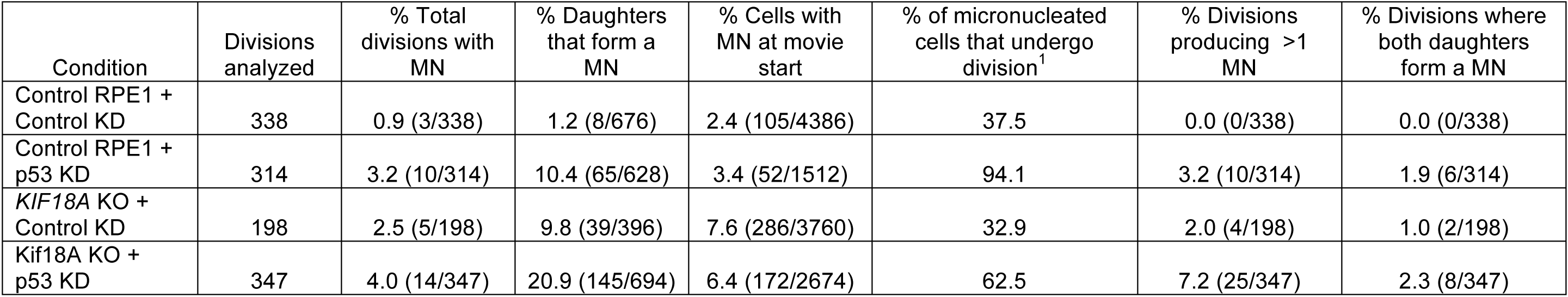
(related to Figures 4 and 7) Summary of live cell analyses of H2B-GFP expressing hTERT-RPE1 cells. Number of divisions analyzed came from at least three independent experiments for each condition. MN = micronucleus. ^1^Calculated at the ratio of percent total divisions with a micronucleus to the percent of cells with a micronucleus at the start of the movie.

